# Disrupted cortex-wide dynamics impair motor planning in Shank3-mutant mice

**DOI:** 10.64898/2026.01.14.699439

**Authors:** Manuel Ambrosone, Elena Montagni, Timothy J. Buschman, Anna Letizia Allegra Mascaro

## Abstract

Coordinated cortical activity underlies voluntary movement planning, yet its disruption is implicated in neurodevelopmental disorders like autism spectrum disorder (ASD). Mutations in the SHANK3 gene, a key ASD risk factor, affect synaptic integrity and cortical network function, but their impact on preparatory cortical dynamics remains unclear. Here, we employed wide-field calcium imaging combined with motion tracking and computational motif analysis to examine cortex-wide activity in Shank3b+/− mice during spontaneous forepaw movements. We found reduced correlation between cortical activity and behavior preceding movement onset, alongside decreased expression of specific cortical motifs involving associative and motor areas. These changes contributed to global functional hyperconnectivity and impaired motor planning. Our findings reveal that disrupted spatiotemporal cortical dynamics in Shank3b+/− mice impair preparatory motor processes, providing insight into neural mechanisms underlying ASD-related motor deficits.

## Introduction

Coordinated cortical activity is essential for the planning and execution of voluntary movements ^1^. During motor behavior, large-scale cortical networks dynamically organize to integrate sensory, associative, and motor information ^2–4^. Notably, cortical activity often precedes movement onset, reflecting preparatory processes that enable the selection and initiation of appropriate motor actions. Such anticipatory activity allows the brain to predict and plan motor consequences before execution ^5–7^. Conversely, disruptions in the temporal coordination between cortical activity and behavior can impair motor planning and adaptability ^8^, deficits frequently observed in neurodevelopmental disorders such as autism spectrum disorder (ASD) ^9,10^.

Among the known genetic causes of ASD, mutations in the SHANK3 gene represent one of the most robust and reproducible monogenic risk factors, making Shank3-mutant mice among the most widely used and well-validated models for investigating the neural and behavioural mechanisms underlying autism ^11^. SHANK3 gene encodes a postsynaptic scaffolding protein critical for the structural and functional integrity of excitatory synapses, and its disruption is causative of Phelan-McDermid syndrome (PMS) ^12–15^. Mice carrying a deletion of the Shank3b isoform recapitulate multiple ASD-relevant phenotypes, including social and cognitive impairments as well as altered motor performance ^16,17^. At the network level, these mice exhibit imbalances in excitatory/inhibitory activity and widespread alterations in functional connectivity (FC) ^18–22^. However, it remains unclear whether and how large-scale deficits affect the predictive dynamics and temporal coordination of cortical networks during movement preparation. Understanding this link is crucial, as preparatory cortical activity provides the neural scaffold for flexible motor control and adaptive behavior ^23,24^.

Recent advances in mesoscopic calcium imaging have made it possible to monitor cortex-wide dynamics with high spatiotemporal precision, enabling direct investigation of how distributed neural activity evolves around behaviorally relevant events ^4,25–30^. This mesoscale approach enables simultaneous monitoring of distributed cortical regions and behaviors with high temporal precision. It also allows the identification of spatiotemporal patterns within a single hemisphere ^31,32^ and across cortex-wide frames ^33–35^. These recurring spatiotemporal patterns of cortical activity, known as ‘motifs’, are characteristic sequences of neural activation underlying relevant behaviors. In mouse models of autism, alterations in cortical dynamics have been observed during transitions between rest and locomotion ^36^, and the motif expression correlates with individual differences in FC and behavior ^37^. A key unresolved question is whether disruptions in motif dynamics are conserved across ASD models or instead reflect model-specific effects of distinct genetic etiologies. Furthermore, the degree to which these disruptions contribute to the observed hyperconnectivity and altered cortex-behavior coupling in Shank3b mice remains unexplored.

Here, we address these gaps by investigating how cortical activity patterns contribute to motor behavior and FC in Shank3b+/− mice using wide-field calcium imaging. By combining motion tracking, convolutional non-negative matrix factorisation (CNMF), and correlation analysis, we examined the relationship between movement timing, cortical motifs, and FC. We found that Shank3b+/− mice exhibit a reduced correlation between cortical activity and behavior in the second preceding movement onset, indicating impaired preparatory dynamics. Moreover, motifs arising from associative and motor cortices show reduced expression in Shank3b+/− mice, contributing both to the loss of predictive cortical activity and to the emergence of global hyperconnectivity. These findings identify a direct link between altered spatiotemporal cortical dynamics and disrupted motor planning in Shank3b+/− mice.

## Results

The aim of this study was to investigate how cortical patterns of activity contribute to motor behavior in Shank3b+/− mice. To this end, during mesoscopic imaging, mice were head-fixed but allowed to move their forepaws freely, in order to directly compare cortical dynamics and global movements on a frame-by-frame basis. At postnatal day 60 (P60), we acquired five recordings of both cortical activity and behavior for each mouse of both genotypes (Supplementary Fig. 1A, N-Shank3b+/+ = 7; N-Shank3b+/− = 9).

### The preparatory phase of the movement is disrupted in Shank3b+/− mice

In order to assess differences in motor behaviors between genotypes, movements were tracked for five different ROIs (eye, snout, whisker, right forepaw, and left forepaw) (Fig. 1A). For each body part, motion energy was computed frame by frame and then averaged. We observed higher motion energy for whiskers and forepaws, while it was almost zero for the eye (Supplementary Fig. 1B) (Eye: 0.73 ± 0.03 AU; Snout: 2.13 ± 0.12 AU; Whisker: 6.94 ± 0.57 AU; R_forepaw: 3.34 ± 0.37 AU; L_forepaw: 2.8 ± 0.38 AU; Nmice = 16; ANOVA One way RM, Tukey correction), indicating that our metric effectively captures differences between large movements (whiskers and forepaws) and relatively fixed body parts (eye and snout). Motion energy values from all body parts were also summed to obtain a single measure of global movement. Comparing average global movement between genotypes revealed no significant differences (Fig. 1B) (Shank3b+/+: 20.0 ± 2.50 AU; Shank3b+/−: 18.8 ± 1.37 AU; p= 0.66, Two sample t test; N-Shank3b+/+ = 7; N-Shank3b+/− = 9).

**Figure 1.**
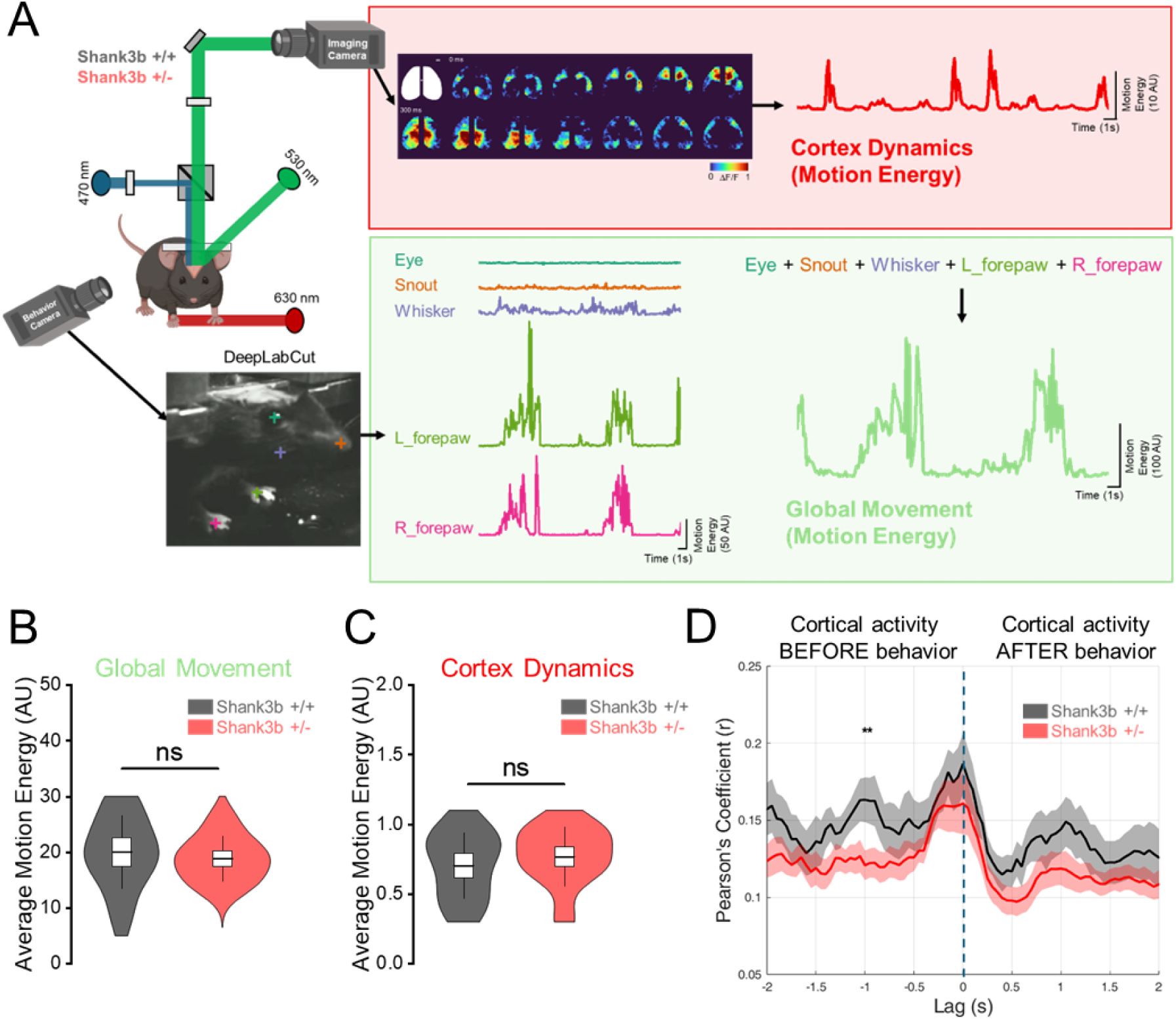
Impaired correlation between cortical dynamics and behavior in Shank3b+/– mice 1 second before movement. (A) Experimental paradigm for tracking cortical activity and behavior in Shank3b+/+ and Shank3b+/– mice. (Top) Cortical dynamics were quantified as the global motion energy of calcium activity in the cortex (red trace), following hemodynamic correction and band-pass filtering (0.4–4 Hz). The white outline indicates the ROI used for cortical activity; the white dot marks the location of bregma; scalebar = 1mm. (Bottom) Global movement was quantified by tracking five ROIs (eye, snout, whisker, right forepaw, left forepaw) using DeepLabCut, and computing the sum of motion energy within these ROIs (green trace). (B-C) Quantification of average motion energy from global movement (B) and cortical dynamics (C) in Shank3b+/+ (black) and Shank3b+/− (red) mice. (D) Correlation between global movement and cortical dynamics at different time lags in Shank3b+/+ (black) and Shank3b+/− (red) mice. The dashed line indicates lag = 0. Asterisks denote statistically significant differences.

To relate behavior to cortical activity, we next quantified the dynamics of cortical activity by computing the motion energy of the calcium imaging data (Fig. 1A). Example traces and montage are shown comparing ΔF/F with cortical motion energy (Supplementary Fig. 1C). While ΔF/F reflects changes in fluorescence intensity, cortical motion energy quantifies the degree of change in cortical activation patterns between consecutive frames: low motion energy indicates similar activity patterns across frames, whereas high motion energy reflects substantial changes in cortical activity. On average, there were no significant differences in cortical motion energy between genotypes (Fig. 1C) (Cortex dynamics-Shank3b+/+: 0.70 ± 0.09 AU; Cortex dynamics-Shank3b+/−: 0.77 ± 0.07 AU; p= 0.57, Two sample t test; N-Shank3b+/+ = 7; N-Shank3b+/− = 9).

Importantly, cortical dynamics and movements were correlated (Supplementary Fig. 1C), highlighting that motor behavior is associated with more dynamic cortical activation. To quantify this relationship across genotypes, we measured the cross-correlation between the motion energy trace of cortical activity and movements for each mouse. Both Shank3b+/+ and Shank3b+/− mice exhibited a correlation between cortical activity and movement at lag 0. Notably, Shank3b+/+ mice showed a stronger correlation 1 second before movement compared to Shank3b+/− mice (Fig. 1D) (Shank3b+/+: 0.16 ± 0.01 r; Shank3b+/−: 0.12 ± 0.01 r; N-Shank3b+/+ = 7; N-Shank3b+/− = 9; *p < 0.05, Two sample t test). These results suggest that a predictive component of cortical activity is disrupted in Shank3b+/− mice.

### Shank3b+/− and Shankb+/+ mice exhibit a similar number of cortical motifs

We hypothesized that this disruption in cortex–behavior coupling could arise from specific cortical signal patterns. To address this, we applied CNMF to identify motifs in cortical activity and assess potential genotype-dependent differences (Fig. 2A). For each mouse, we analyzed four 180-second recordings of resting-state activity, using half of each recording for motif discovery (train epoch) and the other half for evaluation (test epoch) (Supplementary Fig. 1A). From each train epoch, CNMF analysis generated a tensor W representing spatiotemporal motifs over a 1s window, and a matrix H representing their temporal activations. By summing the convolutions between each motif and its temporal activation it is possible to reconstruct recordings (Equation 2) that faithfully reproduced the original data (see representative montages of Fig. 2B). A total of 461 motifs were discovered within the train epochs (Shank3b+/+ = 215; Shank3b+/− = 246), with no significant differences in average motifs’ number between genotypes (Fig. 2C and D) (Shank3b+/+: 8.45 ± 0.61 AU; Nepochs/Nmice= 28/7; Shank3b+/−: 7.50 ± 0.47 AU; Nepochs/Nmice= 36/9; p= 0.22, Two sample t test).

**Figure 2.**
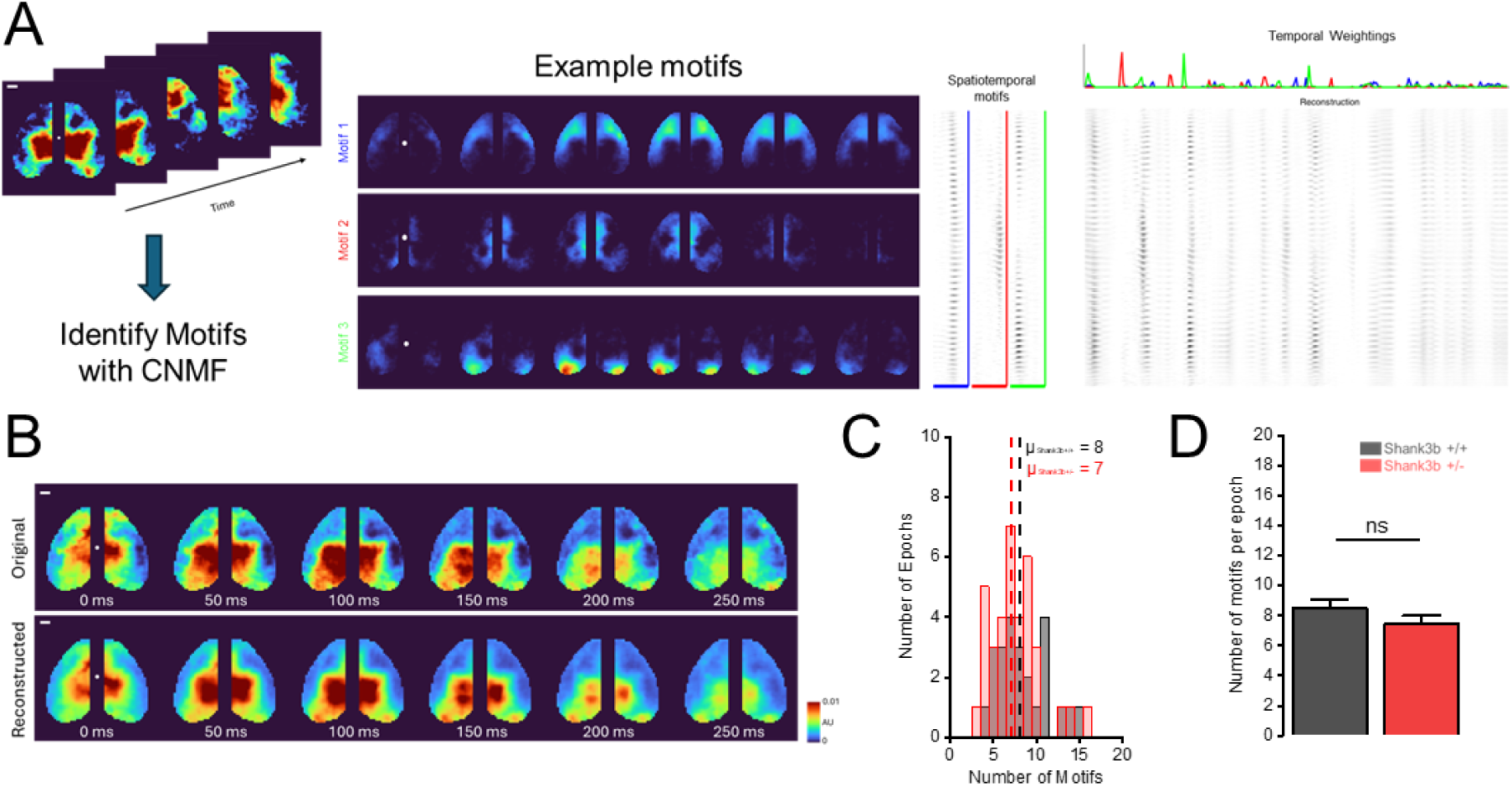
Identification of motifs from cortex-wide activity. (A) Example motifs extracted using CNMF (left), with their corresponding temporal expression profiles (right). The white dot marks the location of bregma; scalebar = 1mm. (B) Comparison between the original cortical activity and the activity reconstructed from the expression of the discovered motifs. The white dot marks the location of bregma; scalebar = 1mm. (C) Quantification of the number of motifs identified in each epoch in each genotype. (D) Average number of discovered motifs per epoch in Shank3b+/+ (black) and Shank3b+/– (red) mice.

To quantify how well the motifs fit neural activity in each animal, we calculated the percentage of variance (PEV) in cortical neural activity that could be explained by the motifs. PEV was measured when refitting to: a) test epochs of the same mouse (within mice PEV) and b) test epochs from different mice (between mice PEV). As expected, there was a marked reduction in PEV when motifs were fit across different individuals (Fig. 3A) (Within mice: 81.6 ± 0.51 %; Between mice-within genotype: 63.7 ± 1.04 %; Between mice-between genotype: 63.5 ± 0.97 %; Nepochs/Nmice = 64/16; ****p < 0.0001; ANOVA One way RM, Tukey correction). This indicates that motifs extracted from a single recording in a given mouse did not fully capture the variance from other mice. However, no significant differences were observed when comparing refitting across mice of the same genotype versus different genotypes, suggesting that this reduction was not genotype-dependent (Fig. 3A) (Between mice-within genotype: 63.7 ± 1.04 %; Between mice-between genotype: 63.5 ± 0.97 %; Nepochs/Nmice = 64/16; p = 0.46; ANOVA One way RM, Tukey correction). Together, these findings show that cortical motifs were preserved across Shank3b+/+ and Shank3b+/− mice, suggesting that the disrupted cortex-behavior coupling may instead relate to how these motifs are expressed or recruited during ongoing cortical activity.

**Figure 3.**
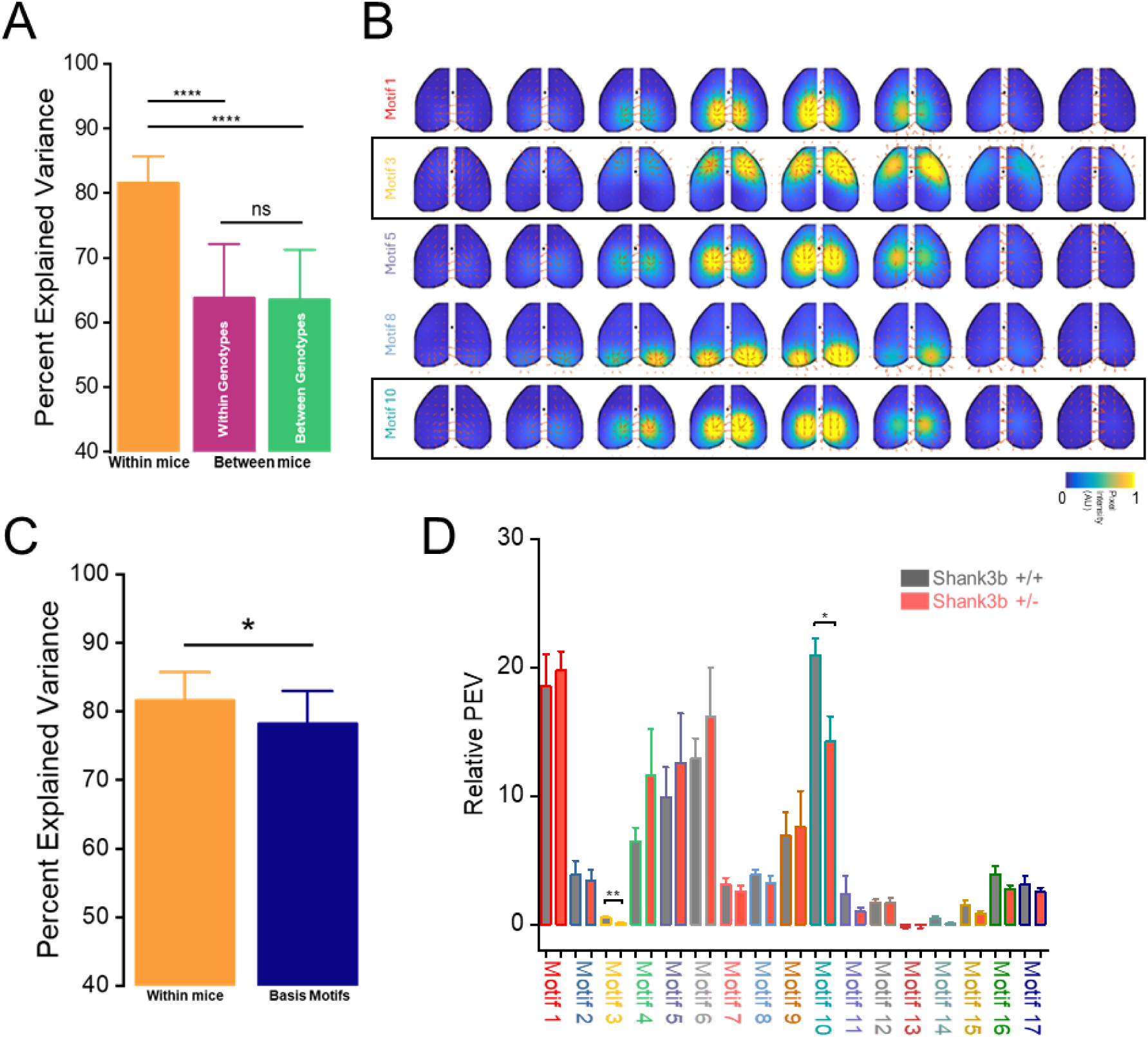
Shank3b+/– mice exhibit altered motifs’ expression. (A) Percent explained variance of the discovered motifs: within individual mice (orange), between mice of the same genotype (purple) and between mice of different genotypes (green). (B) Example basis motifs discovered within genotypes. Black rectangle indicates motifs with different expression. Black dots indicate bregma. (C) Comparison between Percent explained variance of the discovered motifs within individual mice (orange) and for basis motifs (blue). (D) Relative percent explained variance for each basis motif in Shank3b+/+ (black) and Shank3b+/− (red) mice. Asterisks denote statistically significant differences.

### Specific motifs display altered expression between genotypes

To test this, we examined whether specific motifs differed in their expression across mice. To create a common set of motifs that captured canonical spatiotemporal dynamics across all mice and epochs we clustered the identified motifs into a set of ‘basis’ motifs. Clustering was done using an unsupervised algorithm (Phenograph) with the peak of the temporal cross-correlation as the distance metric between two motifs. This resulted in identifying 17 distinct motif clusters (Fig. 3B, Supplementary Fig. 2). The basis motifs reflected the dynamic involvement of one or more brain regions, encompassing both symmetric (e.g., motifs 2, 11, and 14) and asymmetric activation patterns (e.g., motifs 9, 15, and 17). For example, motif 3 captures an activity wave propagating from lateral to medial somatosensory and motor cortices, whereas motif 10 reflects a transient activation confined to parietal and retrosplenial areas. On the other hand, motif 1 and motif 8 represent a localized burst of activity, involving the retrosplenial cortex and visual cortex respectively. Basis motifs captured more of the dynamics than motifs from individual mice, although explained slightly lower PEV compared to fitting motifs within the same mouse (Fig. 3C) (Within mice: 81.6 ± 0.51 %; Basis motifs: 78.2 ± 0.59 %; Nmice = 16; *p < 0.05; Mann-Whitney test). Interestingly, different motifs explained different amounts of variance between genotypes. In particular, motifs 3 and 10 showed significantly reduced PEV in Shank3b+/− mice (Fig. 3D) (Motif 3: Shank3b+/+: 0.52 ± 0.11 %; Shank3b+/−: 0.11 ± 0.04 %; Motif 10: Shank3b+/+: 20.9 ± 1.31 %; Shank3b+/−: 14.2 ± 1.93 %; N-Shank3b+/+ = 7; N-Shank3b+/− = 9; **p < 0.01, *p < 0.05, Two sample t test). Together, these results indicate that motifs 3 and 10 were less expressed in Shank3b+/− mice compared to their non-mutant littermates, reflecting altered spatiotemporal cortical dynamics.

### Reduced motif expression contributes to reduced correlation between cortical activity and behavior in the preparatory phase

We reasoned that specific motifs could contribute to the differences in correlation during the preparatory phase of the movement. To test this, we reconstructed each recording based on individual motif expression and then computed motion energy. We then correlated motion energy for each motif independently with global movement in the second before onset and compared the two genotypes (Fig. 4A). We formulated three hypotheses: (1) different patterns are activated in Shank3b+/+ and Shank3b+/− mice 1 second before movement; (2) the same pattern is activated at different lags in the two genotypes; (3) different patterns are activated at different lags across genotypes. Our analysis revealed that specific motifs (e.g., motifs 5 and 10, arising from associative areas) were preferentially activated 1 second before movement in Shank3b+/+ mice. Each of these motifs showed significantly lower correlation in Shank3b+/− mice at lag –1s (Fig. 4B, Supplementary Fig.3B and C) (Motif 5: Shank3b+/+: 0.10 ± 0.005 r; Shank3b+/−: 0.06 ± 0.005 r; Motif 10: Shank3b+/+: 0.09 ± 0.005 r; Shank3b+/−: 0.06 ± 0.003 r; N-Shank3b+/+ = 7; N-Shank3b+/− = 9; *p < 0.05, Two sample t test). Although not significant, other patterns (e.g., motifs 8 and 9, visual) appeared to be more activated in Shank3b+/− mice at different lags, supporting our third hypothesis (Fig. 4B). Motif 3, involving motor cortices, showed higher correlation in Shank3b+/+ mice 0.6s before movement, while another motor motif (12) peaked at lag –0.3s in Shank3b+/− mice, although this was not significant (Supplementary Fig.3A and D) (Motif 3: Shank3b+/+: 0.08 ± 0.008 r; Shank3b+/−: 0.05 ± 0.002 r; *p < 0.05, Two sample t test; Motif 12: Shank3b+/+: 0.08 ± 0.005 r; Shank3b+/−: 0.12 ± 0.01 r; N-Shank3b+/+ = 7; N-Shank3b+/− = 9; p = 0.06, Two sample t test). Together, these findings suggest that altered recruitment of associative and motor motifs contributes to impaired preparatory cortical dynamics in Shank3b+/− mice.

**Figure 4.**
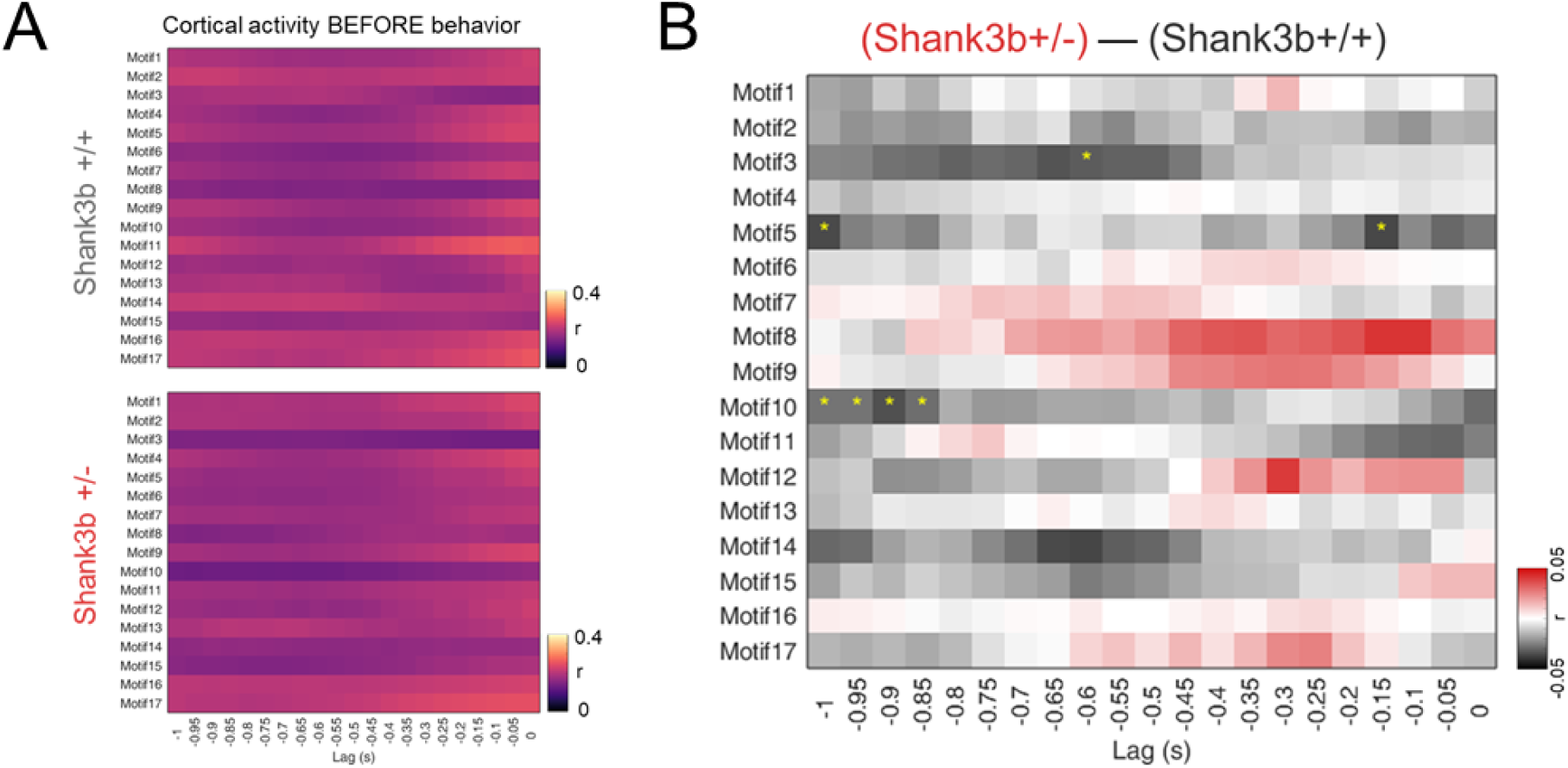
Specific motifs show distinct correlations with global movement. (A) Correlation between global movement and motif dynamics across time lags before behavior in Shank3b+/+ (top) and Shank3b+/− (bottom) mice. (B) Differences in correlation between global movement and motif dynamics across time lags before behavior. Black color indicates higher correlation in Shank3b+/+ mice; red indicates higher correlation in Shank3b+/−mice. Asterisks denote statistically significant differences.

### Specific motifs differentially contribute to hyper-connectivity in Shank3b+/− mice

We next asked whether individual motifs differentially contribute to the large-scale functional architecture of the cortex. We previously demonstrated that Shank3b+/− mice show strong hyper-connectivity at P60 ^22^. By computing FC between areas from reconstructed recordings (baseline) (Fig. 5A) we confirmed the same trend of higher connectivity in Shank3b+/− mice compared to their non-mutant littermates (Fig. 5B and C) (Shank3b+/+ = 0.61 ± 0.02 r; Shank3b+/− = 0.66 ± 0.02 r; N-Shank3b+/+ = 7; N-Shank3b+/− = 9; p= 0.16, Two sample t test). Interestingly, the higher the cortical connectivity, the lower the correlation between cortical activity and behavior at lag –1s (Fig. 5D) (r = −0.24; Nrecordings/Nmice = 64/16; *p < 0.05, Permutation Test). Thus, recordings with higher hyperconnectivity exhibited greater decreases in preparatory activity before movement.

**Figure 5.**
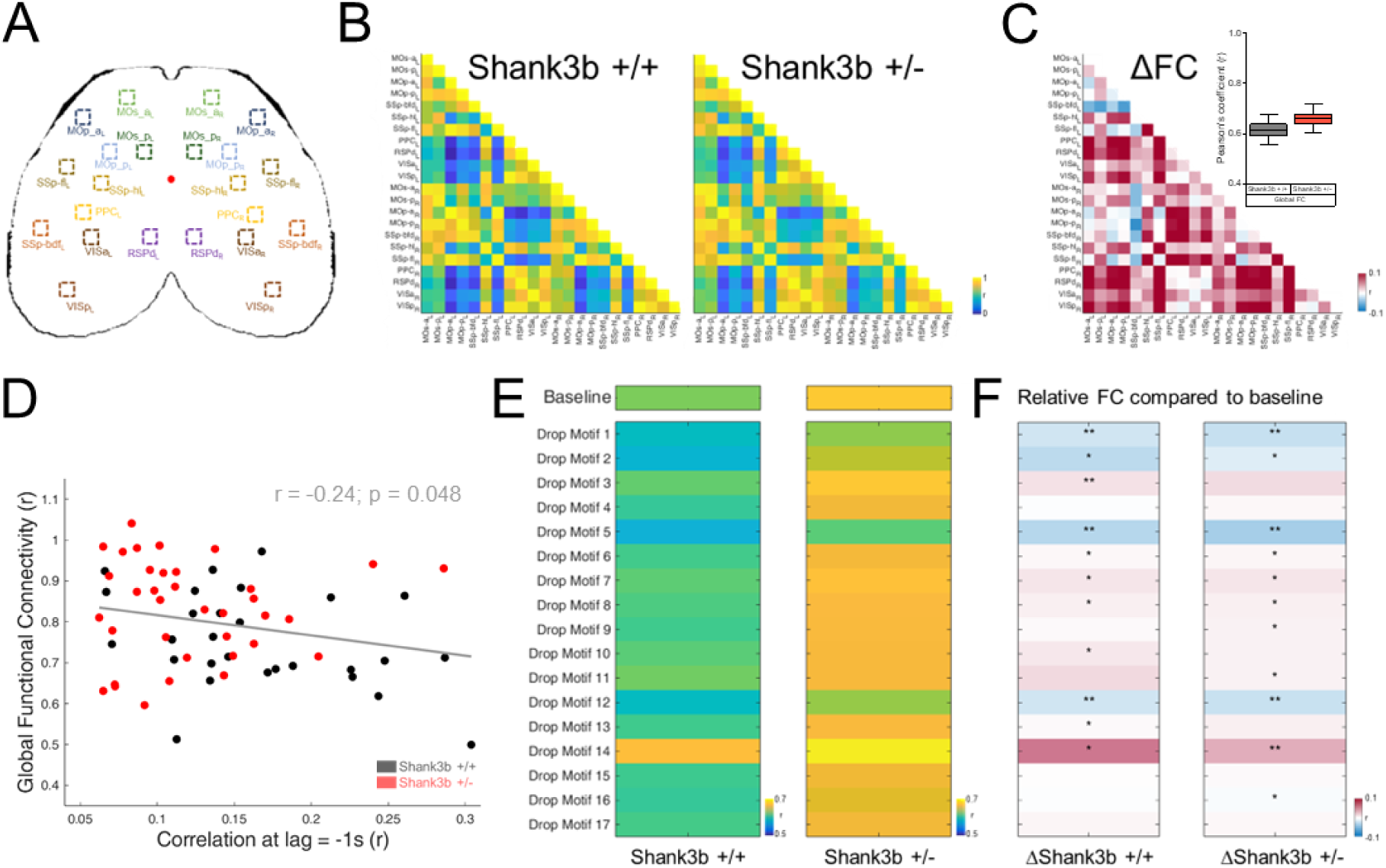
Some motifs contribute more than others to the hyper-connectivity observed in Shank3b+/–mice. (A) Cortical parcellation map indicating selected ROIs across the whole cortex. (B) Functional connectivity (FC) matrices showing Pearson correlation coefficients between cortical areas in Shank3b+/+ (left) and Shank3b+/− mice (right). (C) Difference matrix showing changes in FC between genotypes. Red indicates increased connectivity (hyper-connectivity) in Shank3b+/− mice; blue indicates decreased connectivity (hypo-connectivity). Insert: Quantification of Global FC in Shank3b +/+ (black) and Shank3b+/− (red) mice. (D) Correlation between network level deficit (Higher Global FC) and activity-behavior deficits (Lower correlation at lag = −1) in Shank3b+/+ (black) and Shank3b+/− (red) mice. (E) FC matrices computed after dropping each individual motif, showing global FC in Shank3b +/+ (left), Shank3b+/− (right). (F) Relative FC compared to baseline in Shank3b+/+ (left), Shank3b+/− (right). Red indicates increased connectivity; blue indicates decreased connectivity. Asterisks denote statistically significant differences.

We then systematically removed individual motifs from the total reconstruction to assess their contribution to the hyperconnectivity. Each motif differentially affected FC across genotypes (Fig. 5E). Dropping motifs 6, 7, 8, and 14 increased connectivity in both genotypes, while dropping motifs 1, 2, 5, and 12 induced hypo-connectivity compared to baseline. Removing motifs 3 and 10 (more expressed in Shank3b+/+ mice) increased connectivity only in Shank3b+/+ mice. Conversely, dropping motifs 9, 11, and 16 increased connectivity compared to baseline in Shank3b+/− mice only (see Supplementary Table 2). These findings demonstrate that individual motifs have distinct contributions to the observed FC alterations.

Specifically, our results identify motif 10 as a central contributor to the altered cortical dynamics and functional hyperconnectivity observed in Shank3b+/− mice. Motif 10 is less expressed in Shank3b+/− mice compared to their wild-type littermates and links the observed disruption in correlation between cortical activity and motor activity to aberrant large-scale network organization.

## Discussion

This study reveals that the temporal correlation between cortex-wide signals and behavior, particularly during the preparatory phase of movement, is markedly impaired in Shank3b+/−mice, and that this deficit is linked to a state of network hyperconnectivity.

Motion energy is commonly used for quantifying movement ^38,39^. In our study, we adapted the approach from Hasnain and colleagues’ work ^40^ and combined it with DeepLabCut tracking to obtain a precise frame-by-frame estimate of the movement for different ROIs. As expected, motion energy extracted from forepaws trajectories shows strong temporal fluctuations, whereas the motion energy of the eye remains largely flat, reflecting the distinct dynamics of these body parts (Fig. 1A, Supplementary Fig. 1B). By summing the motion energy across all tracked ROIs, we obtained a robust and sensitive measure of global movement, which captures the overall behavioral output of the animal (Fig. 1B).

To our knowledge, this is the first study in which motion energy is also computed directly from wide-field calcium imaging data to quantify cortical dynamics (Fig. 1C). In Supplementary Fig. 1C, we show that while the average ΔF/F reflects changes in global cortical activity, motion energy captures how much the pattern of cortical activity changes, providing a metric of cortical spatiotemporal dynamics over time. Motion energy remains low when consecutive frames show activation in the same cortical regions, but increases when the set of activated areas changes. In this way, the metric effectively captures rapid changes in cortical activation and offers a robust measure of how cortical activity patterns vary across consecutive imaging frames.

The key functional deficit we identified is a reduction in the ability of cortical activity to predict movement in Shank3b+/− mice, expressed as a reduced correlation between cortical signals and movement 1 second before the onset (Fig. 1D). This temporal window corresponds to the period of motor planning and readiness ^7^. In both rodents and primates, self-initiated actions are preceded by slow, widespread cortical ramping activity that reflects the transition into the “movement-potent” neural subspace required for action initiation ^29,40,41^. The reduction in predictive correlation therefore suggests that Shank3 haploinsufficiency disrupts the emergence, stability, or coordination of this preparatory state. This is in line with previous studies showing impaired motor activity in Shank3 mutant mice with marked impairments in motor coordination and endurance ^17,42,43^.

To uncover which spatiotemporal patterns contribute to this deficit, we extracted low-dimensional motifs from calcium data (Fig. 2) ^33,44^. Phenograph clustering identified 17 basis motifs, some of which appeared partially similar to one another (Supplementary Fig. 2). For example, motifs 1 and 4 both involve the retrosplenial cortices, while motifs 6 and 9 are predominantly expressed in visual cortices. However, they show marked differences in expression (Fig. 3D), suggesting that these motifs play divergent roles in shaping cortical dynamics between genotypes despite their spatial similarity. Notably, similar motifs disruptions were described in the valproic acid mouse model of autism ^37^, indicating that abnormal expression of mesoscale cortical primitives may be a convergent hallmark for ASD.

Two motifs were particularly relevant to explain the differences between the two genotypes (Fig 3D). First, motif 3 captures a propagating sequence across somatosensory and motor cortices, regions crucial for integrating sensory information to guide behavior ^34^. Its reduced expression in Shank3b+/− mice aligns with evidence that SHANK3 deletion impairs motor processing, as demonstrated by poor performance on the rotarod and grid hanging tests ^43^. Second, motif 10 predominantly involves parietal and retrosplenial cortices, key associative hubs ^45^. They play essential roles in integrating spatial, and motor-related information, supporting top-down control of motor planning and spatially guided actions ^46–48^. Given that SHANK3 mutations disrupt long-range prefrontal and associative connectivity ^15,19,22^, its reduced expression likely reflects a dysfunction in high-level integrative networks.

We then directly assessed whether these motifs contribute to the impaired preparatory timing. In wild-type mice, associative motifs (5 and 10) were preferentially activated at −1 s relative to movement onset (Fig. 4; Supplementary Fig. 3), consistent with their role in high-order motor planning ^49–51^. In contrast, Shank3b+/− mice showed significantly lower correlations at this critical preparatory period. This difference suggests a disrupted transition from associative preparation to motor execution. Similar disruptions of transitions in associative cortical dynamics have been reported in other ASD models, such as the 15q11-13 duplication line ^36^.

To analyze the network organization underlying these timing abnormalities, we examined the functional connectivity across cortical areas. We identified a negative correlation between global FC and the cortex-behavior impairment at −1 s (Fig. 5D). Thus, mice exhibiting the highest levels of functional hyperconnectivity exhibit the most severe impairments in preparatory timing. SHANK3 deletion induces early corticostriatal hyperactivity and premature circuit maturation ^18^, leading to E/I imbalance and altered long-range coupling ^20^. We propose that this increased network strength constrains the system’s ability to transiently segregate or reconfigure activity patterns during behavioral transitions.

The motif-specific FC analysis supports this idea: removing motifs 3 and 10 increased FC only in control animals (Fig. 5E, Supplementary Table 2), possibly indicating that the expression of these motifs is important in maintaining lower connectivity levels. Their diminished expression in mutants, therefore, contributes directly to the hyperconnected baseline. Notably, motif 10 emerges as a central regulator, tuning the global FC to orchestrate the correct pattern for cortical motor planning.

Together, these results identify motif 10 as a key regulator of balanced mesoscale dynamics in the preparatory phase of the behavior in the healthy cortex. Its systematic disruption in Shank3b-haploinsufficient mice establishes a direct relationship between impaired motifs’ expression, aberrant network function, and deficits in self-generated action. Taking advantage of modern optogenetic tools capable of temporally precise excitation or inhibition of cortical activity ^52–55^, future work can expand toward closed-loop interventions to modulate, and eventually restore, motif-level dynamics in neurodevelopmental disorders.

## Acknowledgements

This work has been funded by Telethon Seed Grants GSA22E006 and GSA25E002 (ALAM), the Italian Ministry of Universities and Research PRIN 2022 2022YCTLPL (ALAM), project THE Tuscany Health Ecosystem ECS_00000017 MUR_ PNRR (ALAM), and NIMH R01MH126022 (TJB). The project was done, in part, while MA was a visiting scholar at Princeton Neuroscience Institute.

## Author Contributions (CRediT)

MA and ALAM conceived the study.

MA, ALAM and TJB designed and interpreted experiments and computational analyses.

MA and EM performed experiments and generated data.

MA and EM conducted computational analyses.

MA and ALAM drafted the manuscript.

TJB edited the manuscript.

ALAM and TJB contributed funding and resources.

All authors participated in the review of the manuscript.

## Data and Code Availability Statement

The datasets and code generated during this study are available from the corresponding author upon reasonable request.

## Declaration of generative AI and AI-assisted technologies in the manuscript preparation process

During the preparation of this work the authors used ChatGPT in order to improve the syntax and the readability of the manuscript. After using this tool/service, the authors reviewed and edited the content as needed and take full responsibility for the content of the published article.

## Materials and Methods

### Ethical statement

A total of 16 B6.129-Shank3^tm2Gfng^/J mice (2 months old; 9 females and 7 males) were used in the study. Animals were housed in standard cages under a 12 h light/dark cycle with food and water provided ad libitum. All experimental procedures complied with relevant guidelines and regulations and were approved by the Italian Ministry of Health (Authorization n. 721/2020). Mice were assigned to experimental groups based on genotype, and investigators were blinded to group allocation during both experiments and behavioral analyses.

### Virus injection for the expression of GCaMP7f

To achieve selective expression of the genetically encoded calcium indicator GCaMP7f in excitatory neurons, we systemically delivered a viral vector (ssAAV-PHP.eB/2-mCaMKIIα-jGCaMP7f-WPRE-bGHp(A); titer: 1.3 × 10¹³ vg/mL; Viral Vector Facility, CH) via the retro-orbital sinus. Injections were performed on postnatal day 30 (P30) under isoflurane anesthesia. Viral stocks were diluted in sterile saline to a final volume of 150 µL.

### Intact skull window surgery

One week after viral delivery, mice underwent preparatory surgery for chronic imaging under head fixation. Anesthesia was induced with 3% isoflurane and maintained at 1–2%, and animals were positioned in a stereotaxic frame (KOPF, model 1900). Ophthalmic gel (Lacrilube) was applied to prevent corneal desiccation, and body temperature was maintained at 37 °C using a feedback-controlled heating pad. A local anesthetic (2% lidocaine) was applied to the scalp before making a midline incision. Skin and periosteum were carefully removed, and cranial landmarks (bregma and lambda) were identified and marked. A custom aluminum head bar was secured posterior to lambda with dental cement (Super Bond C&B, Sun Medical), and the remaining exposed skull was sealed with transparent cement to create a stable preparation for imaging through the intact skull.

### Wide-field microscope

Wide-field imaging was performed using a custom-made microscope (see Conti et al., 2019) ^56^. GCaMP7f excitation was achieved with a 470 nm LED (M470L3, Thorlabs) filtered through a 482/18 nm bandpass (Semrock) and reflected onto the cortical surface via a dichroic mirror (DC FF 495-DI02, Semrock) through a 2X Super Apochromatic objective (TL2X-SAP, 0.1 NA, 56.3 mm working distance; Thorlabs).

Reflectance signals were acquired using a 530 nm LED (M530L4, Thorlabs) positioned at a 45° angle relative to the cortex. Excitation and reflectance channels were alternated at 20 Hz using stroboscopic illumination. Emitted fluorescence and reflectance were filtered through a 525/50 nm bandpass (Semrock) and recorded with a CMOS camera (ORCA-Flash4.0 V3, C13440-20CU; Hamamatsu) at 40 Hz, with a spatial resolution of 512 × 512 pixels, corresponding to a field of view of ∼11.5 × 11.5 mm.

### Awake imaging and behavior acquisition

Mice were gradually habituated to head fixation over two consecutive days, 15 minutes per day, to reduce stress and minimize sudden movements during recordings. Imaging sessions were conducted at postnatal day 60. Imaging stacks were aligned using custom software, which centered the field of view (FOV) by taking into account the bregma and lambda landmarks previously marked on the skull. This approach enabled precise alignment of imaging sessions across animals and time points using anatomical references. Each session comprised five 180-second recordings capturing spontaneous cortical activity in awake, resting mice. Animals were head-fixed but free to move their limbs and were not performing any specific task.

To monitor movement during imaging, a camera (PointGrey FLIR Chameleon3, CM3-U3-13Y3C-CS) was placed 10 cm in front of the mouse, recording at 40 frames per second with 512 × 512 pixel resolution to cover the frontal area. A 630 nm visible light source illuminated the forepaws to facilitate movement detection.

### Data analysis

All data analyses were performed in MATLAB (MathWorks), ImageJ, Python, DeepLabCut, and Origin.

#### Movements tracking using DeepLabCut

At P60, behavioral videos were analyzed using Python. Alternate frames from each video were taken for the alignment to the imaging recording. The resulting videos were then preprocessed to increase the contrast. Five regions of interest (Eye, Snout, Whisker, Right forepaw and Left forepaw) were tracked with DeepLabCut extracting the spatial (x, y) coordinates for each frame ^57^. The first recording was used as habituation and to train DeepLabCut to extract ROI coordinates for each frame (Supplementary Fig. 1A).

#### Motion energy of global movements

Adapted from Hasnain and colleagues ^40^, the motion energy for a given frame and ROI was defined as the absolute value of the difference between the value of the coordinates of that frame and the next frame. The global movement for each frame was then defined as the sum of motion energy values across all selected ROIs, yielding a global index of motor activity. This approach is sensitive to rapid, localized movements (e.g., forepaws) while remaining relatively unaffected by slow and widespread changes such as those associated with respiration.

#### Image preprocessing

Image stacks collected across sessions for each mouse were aligned using custom software based on the anatomical landmarks bregma and lambda. A subject-specific FOV template was employed to manually fine-tune the imaging area on each recording day. To assign functional signals to distinct cortical regions, the imaged cortex was registered to a two-dimensional projection of the Allen Mouse Brain Atlas (www.brain-map.org), aligned to the imaging plane. For each recording block, ΔF/F₀ was computed at the pixel level, where ΔF represents the change in fluorescence intensity relative to the mean fluorescence (F₀) across the session. Hemodynamic correction was applied using the ratiometric method described by Scott and colleagues ^58^:

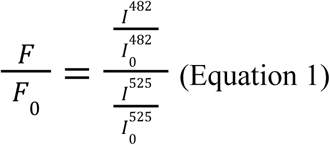

where I^482^ and I^525^ correspond to the fluorescence and reflectance signals at 482 nm and 525 nm, respectively.

Following ΔF/F₀ calculation and hemodynamic correction, images were downsampled to 64 × 64 pixels and registered to a 2D projection of the Allen Brain Atlas using bregma and lambda coordinates. The recordings were then band-pass filtered between 0.4 and 4 Hz ^59,60^ and masked to isolate the cortical area of interest.

#### Motion energy of cortical activity

In order to quantify the global cortical dynamic from the preprocessed images, the motion energy for a given frame and pixel was computed as the absolute value of the difference between consecutive frames, resulting in a matrix of size pixels × time points. The cortex dynamic for each frame was then calculated by summing the motion energy values across all pixels for that frame and then squaring the result.

#### Cross-correlation between cortical dynamics and global movements

For each recording, the resulting motion energy traces of movements and cortical activity were aligned. Pearson’s correlation coefficients were calculated at multiple time lags and then averaged within recordings and genotypes.

#### Motifs discovery

Spatiotemporal sequences in widefield imaging data were identified using the seqNMF algorithm ^44^, which applies CNMF with an additional regularization term to promote the discovery of repeating patterns ^33^.

To obtain a more accurate estimation of the underlying neural activity, we performed pixel-wise deconvolution of the ΔF/F signal using the Lucy-Richardson deconvolution algorithm after image preprocessing (implemented via the “lucid” function; see Stern et al., 2020 ^61^, with parameters gamma = 0.95, smt = 1, and p_num = 30). Following deconvolution, pixel values were normalized to a range between 0 and 1. For each mouse, four recordings were divided into two 90-second epochs; alternating chunks were used for motif discovery and testing. Each chunk was vectorized into a P × 1 vector (P = number of pixels), and each recording epoch was represented as a P × T matrix (T = number of time points). This matrix was factorized into K motifs of size P × L, forming a tensor W (P × K × L) representing short spatiotemporal motifs of images (P) over short durations of time (L time points), and a matrix H (K × T) representing their temporal activations. The original data were approximated by the sum of convolutions between each motif and its temporal activation:

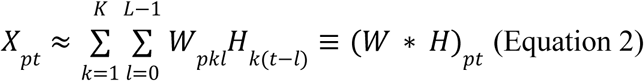

The matrices W and H were optimized using a multiplicative update algorithm. Motif duration (L = 20 frames, 1 s) and number of motifs (K = 28) were chosen to capture all repeating sequences without constraining the solution ^33^.

To reduce redundancy, seqNMF includes a spatiotemporal penalty that discourages multiple motifs from describing the same sequence, prevents motif splitting across time, and promotes temporal independence. The parameter λ (set to 0.0005) balances reconstruction accuracy, motif separation, and the number of motifs discovered. To encourage event-based motifs and penalize overlap in the temporal activations, an additional cost term (λ_*Hortho*_) was implemented and set to 1. All factorizations were run for 300 iterations, which was sufficient for convergence of the cost function as reported by MacDowell and Buschman ^33^. A complete list of the seqNMF parameters used is provided in Supplementary Table 1.

#### Refitting Motifs to Withheld Data

To assess motif expression, motifs identified during the discovery phase were refitted to withheld recording chunks using the seqNMF algorithm. During refitting, the spatial components W (i.e., the basis motifs) were held constant, and only the temporal activations H were updated. This procedure allowed adjustment of motif timing without altering their spatial structure. Because only the temporal coefficients were optimized, the regularization parameter λ was set to zero. Unlike the discovery phase, which included penalties to enforce spatial and temporal independence among motifs, refitted motifs were not constrained by such regularization. As a result, some combinatorial activation of motifs was observed. All parameters used for refitting are listed in Supplementary Table 1.

#### Generating Basis Motifs

Basis motifs were identified using the Phenograph algorithm ^33,62^. Motifs were first rescaled to [0,1] and smoothed with a 3D Gaussian filter (σ = [1,1,0.1]). Each motif was connected to its *k* (here, 17) nearest neighbors based on peak temporal cross-correlation, and clusters were defined via Louvain community detection. Within each cluster, the top 10% of motifs with the most within-cluster neighbors were designated as the core set. Core motifs were aligned to a reference template (motif with highest zero-lag correlations) and zero-padded to 3L (60 time points) to account for temporal offsets. Finally, the basis motifs were aligned to one another by centering their temporal activity around the midpoint. Time points with zero variance across motifs were excluded, yielding final basis motifs of up to 40 time points (∼2 s). Detailed descriptions are provided in Supplementary Table 1.

#### Evaluating motifs expression

Percent explained variance (PEV) was used to quantify how well reconstructed data captured the original neural activity:

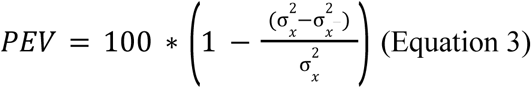

where 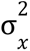 and represents the total spatiotemporal variance of the original data, and 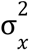–is the variance of the data reconstructed from the factorization.

To assess individual motif contributions, each motif was convolved with its temporal activation and PEV was computed. This measure reflects both the frequency of motif occurrence and its fidelity to underlying neural dynamics. Relative contribution of each motif was obtained by dividing the PEV of that motif alone by the total PEV explained by all motifs in the epoch.

#### Cross-correlation between motif dynamics and global movements

The seqNMF algorithm also allows reconstruction of data based on individual motif expressions. For each motif, motion energy was computed from the reconstructed recording using the same procedure as for the original data and then aligned to the global movement motion energy. Pearson’s correlation coefficients were then calculated for each motif across different time lags (from −20 to 0) and averaged within recordings and across genotypes. To evaluate genotype differences, contrast matrices were created by subtracting the mean r-values of Shank3b+/+ mice from those of heterozygous Shank3b+/− mice.

#### Cortical Parcellation

A total of 22 ROIs were selected from reconstructed images (11 ROI for each hemisphere, 4×4 pixels). The abbreviations and extended names for each areas are as follows: MOs-a, anterior region of secondary motor cortex; MOs-p, posterior region of secondary motor cortex; MOp-a, anterior region of primary motor cortex; MOp-p, posterior region of primary motor cortex; SSp-bfd, primary somatosensory area, barrefield; SSp-fL, primary somatosensory area, forelimb; SSp-hl, primary somatosensory area, hindlimb; PPC, posterior parietal cortex; RSPd, retrosplenial cortex; VISa, associative visual cortex; VISp, primary visual cortex. Suffixes L and R were added to denote cortical areas of the left or right hemisphere, respectively (e.g., MOp-a_L_, MOp-a_R_).

#### Resting state functional connectivity

FC was computed for each recording by calculating Pearson’s correlation coefficients between ROIs. Correlations were estimated from the average fluorescence signal across the spatial extent of each ROI within a 180-second time window. Correlation values were then averaged across imaging sessions. To enable group-level comparisons, individual correlation matrices were transformed using Fisher’s r-to-z transformation and subsequently averaged across animals within each experimental group. The resulting group-averaged matrices were then converted back to correlation coefficients (r-values).

To assess genotype differences, we generated contrast matrices by subtracting the mean r-values of Shank3b+/+ mice from those of heterozygous Shank3b+/− mice for each ROI pair.

For motif-specific FC, connectivity was computed by systematically removing each motif from the full reconstruction (baseline) and recalculating the correlations. Mean r-values for each ROI pair were then averaged across areas to obtain a single FC value for each recording and motif.

### Statistical analysis

Two sample t-tests were used to quantify differences between groups in cortical dynamics, global movements, correlation between cortical dynamics and movements, number of motifs, motifs expression, and global FC. One-way repeated measures (RM) ANOVA followed by Tukey correction was used to investigate differences in average motion energy between body parts and differences between PEV within and between mice. Mann-Whitney test was used for comparison between PEV within mice and basis motifs. A permutation test was used to quantify the correlation between global FC and correlation at lag = −1s (Number of permutations = 1000). Errors are reported as Standard Error of Means, *p < 0.05, ** p < 0.01, *** p < 0.001, **** p<0.0001.

---

**Supplementary Figure 1.**
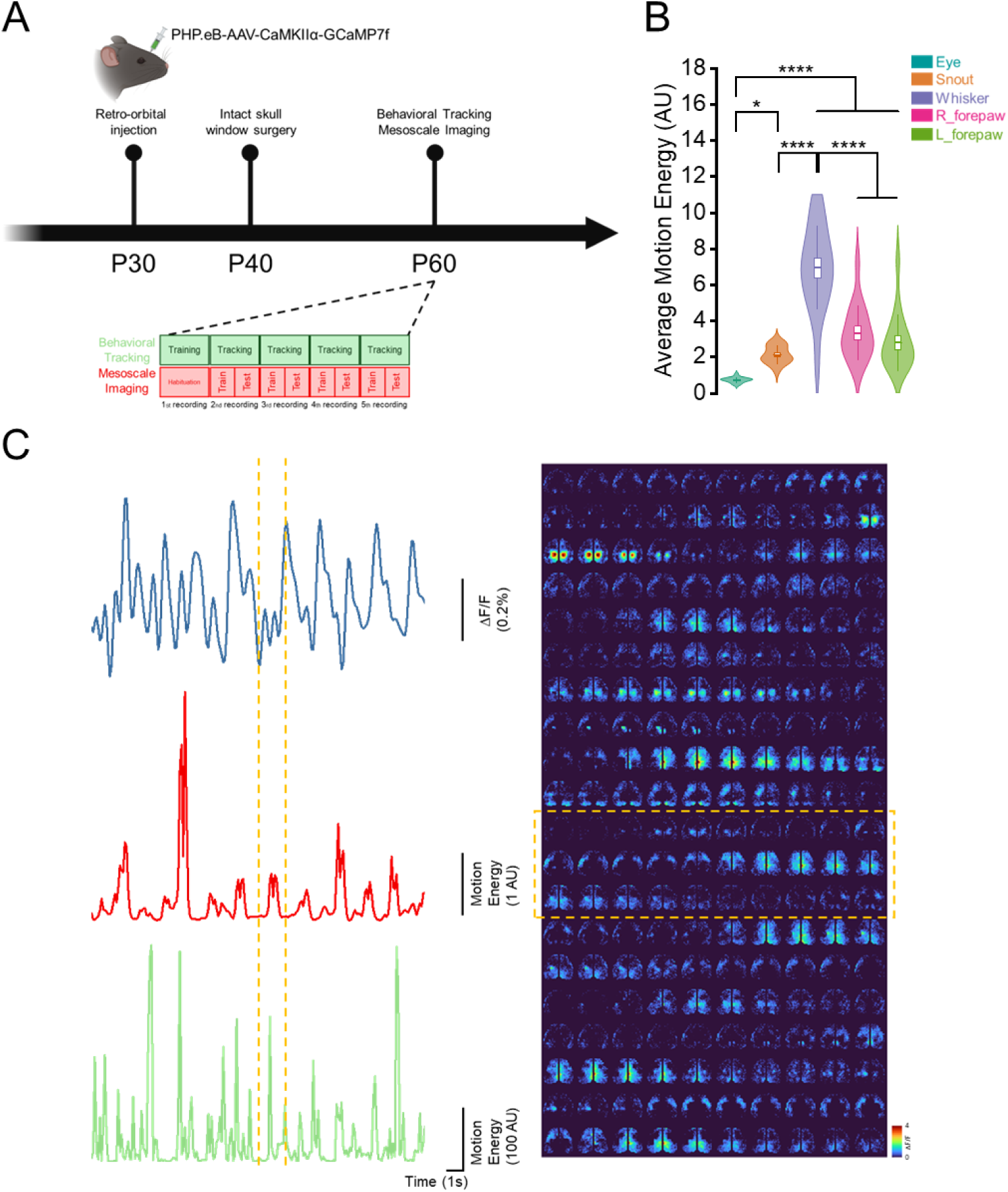
Experimental timeline and motion energy calculation. (A) Experimental timeline showing retro-orbital injection on PHP.eB-AAV at P30, followed by Intact skull window surgery at P40 and Imaging at P60. Five recordings were collected for each mouse. The first one was used to train DeepLabCut. (B) Quantification of average motion energy for each ROI (eye snout, whisker, right forepaw, left forepaw). Asterisks denote significant differences. (C) (Left) Example traces showing the comparison between ΔF/F (top), Cortex dynamics (middle) and Global Movements (bottom) through time. (Right) Example montage of cortical activity through time.

**Supplementary Figure 2.**
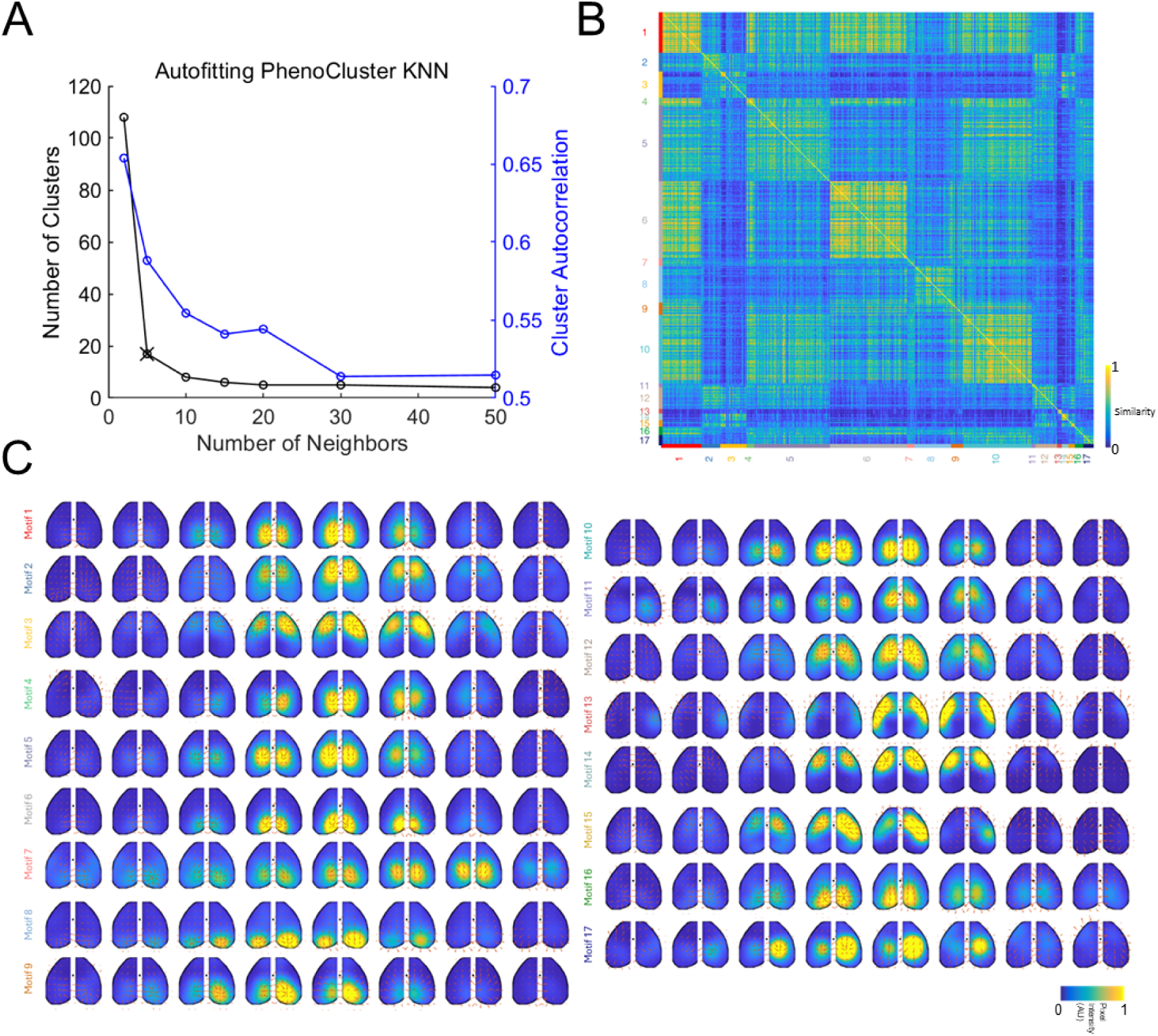
Cluster parameters. (A) Autofitting PhenoCluster KNN showing the optimal number of clusters for basis motifs (black cross, K=17). (B) Pairwise correlation matrix showing the similarity across all motifs. (C) Representation of all basis motifs extracted from discovered motifs.

**Supplementary Figure 3.**
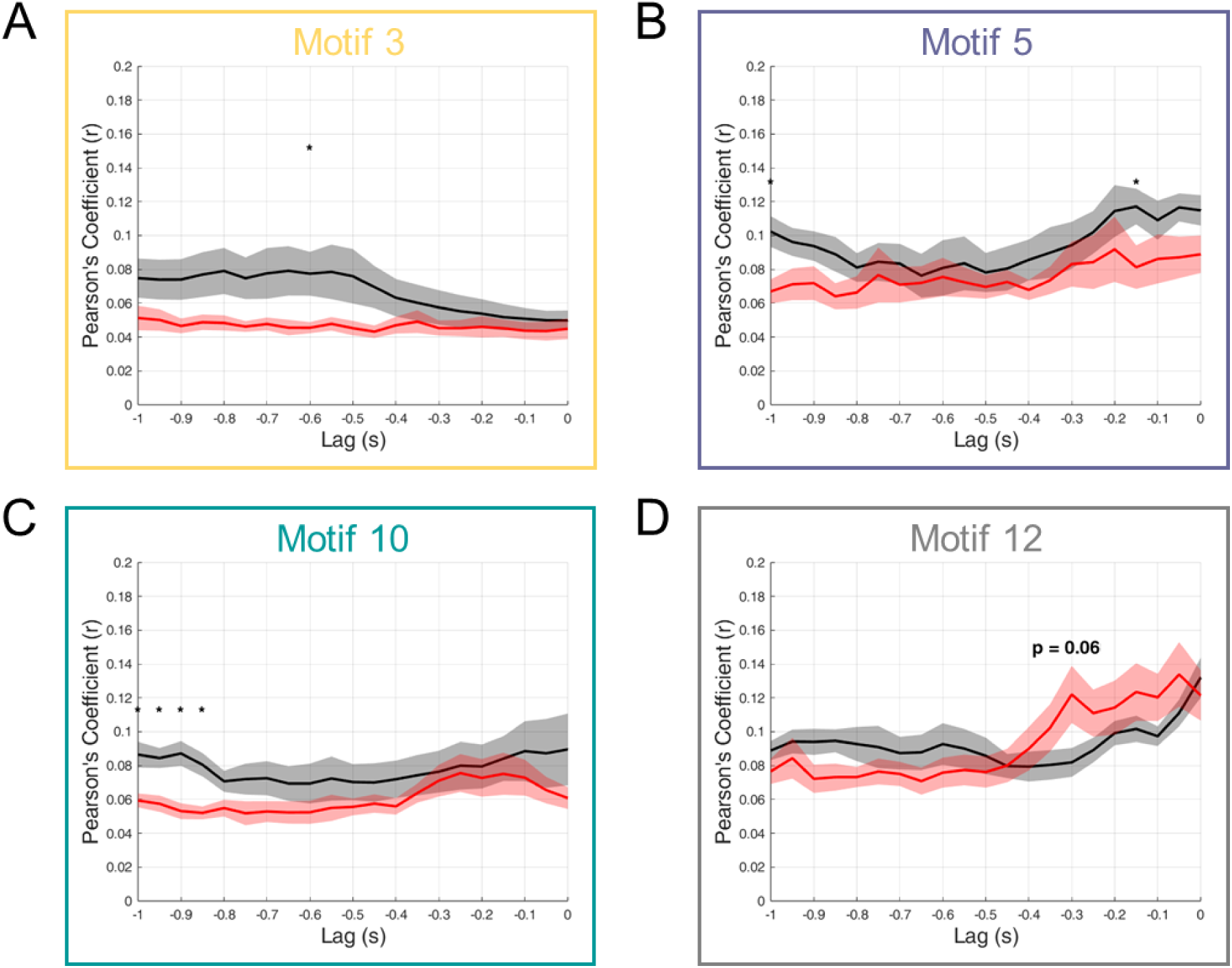
Correlation between specific motifs dynamic and global movement. (A-B) Correlation between global movement and cortical dynamics at negative lags for motif 3 (A), motif5 (B), motif 10 (C) and motif 12 (D) in Shank3b+/+ (black) and Shank3b+/– (red) mice. Asterisks denote statistically significant differences.

**Supplementary Table 1.** CNMF parameters used for motif discovery and refit.

|  | Motifs Discovery | Motifs Refit |
| --- | --- | --- |
| K | 28 | 17 (# basis motifs) |
| L | 20 | 20 |
| $\lambda$ | 0.0005 | 0 |
| W | Random | Basis Motifs |
| H | Random | Random |
| $\lambda_{Hortho}$ | 1 | 1 |
| $\lambda_{Wortho}$ | 0 | 0 |
| Iterations | 300 | 100 |
| $W_{fixed}$ | 0 | 1 |
| $W_{updated}$ | 1 | 0 |

**Supplementary Table 2.** FC average values and SEM in Shank3b+/+ (black) and Shank3b+/− (red) mice. P values indicate significance when comparing each value with the value of the baseline for each group.

|  | FC<br>Shank3b+/+<br>(r) | SEM<br>Shank3b+/+<br>(r) | P value | FC<br>Shank3b+/-<br>(r) | SEM<br>Shank3b+/-<br>(r) | P value |
| --- | --- | --- | --- | --- | --- | --- |
| Baseline | 0.615 | 0.013 | // | 0.659 | 0.011 | // |
| Drop motif 1 | 0.596 | 0.013 | 0.001 | 0.637 | 0.011 | 0.0001 |
| Drop motif 2 | 0.589 | 0.014 | 0.022 | 0.647 | 0.012 | 0.044 |
| Drop motif 3 | 0.627 | 0.012 | 0.004 | 0.674 | 0.01 | 0.057 |
| Drop motif 4 | 0.614 | 0.013 | 0.182 | 0.663 | 0.011 | 0.412 |
| Drop motif 5 | 0.586 | 0.014 | 0.002 | 0.624 | 0.012 | 0.004 |
| Drop motif 6 | 0.619 | 0.013 | 0.031 | 0.664 | 0.011 | 0.046 |
| Drop motif 7 | 0.626 | 0.012 | 0.019 | 0.67 | 0.011 | 0.026 |
| Drop motif 8 | 0.621 | 0.013 | 0.035 | 0.666 | 0.011 | 0.033 |
| Drop motif 9 | 0.617 | 0.012 | 0.066 | 0.665 | 0.011 | 0.043 |
| Drop motif 10 | 0.624 | 0.012 | 0.023 | 0.665 | 0.011 | 0.228 |
| Drop motif 11 | 0.63 | 0.012 | 0.155 | 0.665 | 0.011 | 0.018 |
| Drop motif 12 | 0.597 | 0.013 | 0.0001 | 0.638 | 0.012 | 0.0002 |
| Drop motif 13 | 0.618 | 0.013 | 0.049 | 0.666 | 0.011 | 0.209 |
| Drop motif 14 | 0.669 | 0.011 | 0.015 | 0.692 | 0.01 | 0.001 |
| Drop motif 15 | 0.616 | 0.013 | 0.829 | 0.662 | 0.011 | 0.431 |
| Drop motif 16 | 0.614 | 0.013 | 0.595 | 0.657 | 0.011 | 0.013 |
| Drop motif 17 | 0.619 | 0.013 | 0.067 | 0.663 | 0.011 | 0.148 |

## References

1. Morandell, K., and Huber, D. (2017). The role of forelimb motor cortex areas in goal directed action in mice. Sci. Rep. 7, 15759.

2. Ferezou, I., Haiss, F., Gentet, L.J., Aronoff, R., Weber, B., and Petersen, C.C.H. (2007). Spatiotemporal dynamics of cortical sensorimotor integration in behaving mice. Neuron 56, 907–923.

3. Musall, S., Kaufman, M.T., Juavinett, A.L., Gluf, S., and Churchland, A.K. (2019). Single-trial neural dynamics are dominated by richly varied movements. Nat. Neurosci. 22, 1677–1686.

4. Cardin, J.A., Crair, M.C., and Higley, M.J. (2020). Mesoscopic imaging: Shining a wide light on large-scale neural dynamics. Neuron 108, 33–43.

5. Weinrich, M., Wise, S.P., and Mauritz, K.H. (1984). A neurophysiological study of the premotor cortex in the rhesus monkey. Brain 107 (Pt 2), 385–414.

6. Churchland, M.M., Cunningham, J.P., Kaufman, M.T., Ryu, S.I., and Shenoy, K.V. (2010). Cortical preparatory activity: representation of movement or first cog in a dynamical machine? Neuron 68, 387–400.

7. Mitelut, C., Zhang, Y., Sekino, Y., Boyd, J.D., Bollanos, F., Swindale, N.V., Silasi, G., Saxena, S., and Murphy, T.H. (2022). Mesoscale cortex-wide neural dynamics predict self-initiated actions in mice several seconds prior to movement. Elife 11, e76506.

8. Inagaki, H.K., Chen, S., Ridder, M.C., Sah, P., Li, N., Yang, Z., Hasanbegovic, H., Gao, Z., Gerfen, C.R., and Svoboda, K. (2022). A midbrain-thalamus-cortex circuit reorganizes cortical dynamics to initiate movement. Cell 185, 1065–1081.e23.

9. Mahajan, R., Dirlikov, B., Crocetti, D., and Mostofsky, S.H. (2016). Motor circuit anatomy in children with autism spectrum disorder with or without attention deficit hyperactivity disorder: Motor circuit anatomy in ASD with or without ADHD. Autism Res. 9, 67–81.

10. Yoo, G.E., Lee, E.Y., and Lee, E. (2025). Neural correlates of social motor coordination in autism: A systematic review and meta-analysis of fNIRS studies. Neurosci. Biobehav. Rev. 177, 106347.

11. Monteiro, P., and Feng, G. (2017). SHANK proteins: roles at the synapse and in autism spectrum disorder. Nat. Rev. Neurosci. 18, 147–157.

12. Bonaglia, M.C., Giorda, R., Borgatti, R., Felisari, G., Gagliardi, C., Selicorni, A., and Zuffardi, O. (2001). Disruption of the ProSAP2 gene in a t(12;22)(q24.1;q13.3) is associated with the 22q13.3 deletion syndrome. Am. J. Hum. Genet. 69, 261–268.

13. Durand, C.M., Perroy, J., Loll, F., Perrais, D., Fagni, L., Bourgeron, T., Montcouquiol, M., and Sans, N. (2012). SHANK3 mutations identified in autism lead to modification of dendritic spine morphology via an actin-dependent mechanism. Mol. Psychiatry 17, 71–84.

14. Costales, J.L., and Kolevzon, A. (2015). Phelan-McDermid syndrome and SHANK3: Implications for treatment. Neurotherapeutics 12, 620–630.

15. Delling, J.P., and Boeckers, T.M. (2021). Comparison of SHANK3 deficiency in animal models: phenotypes, treatment strategies, and translational implications. J. Neurodev. Disord. 13, 55.

16. Peça, J., Feliciano, C., Ting, J.T., Wang, W., Wells, M.F., Venkatraman, T.N., Lascola, C.D., Fu, Z., and Feng, G. (2011). Shank3 mutant mice display autistic-like behaviours and striatal dysfunction. Nature 472, 437–442.

17. Wang, X., McCoy, P.A., Rodriguiz, R.M., Pan, Y., Je, H.S., Roberts, A.C., Kim, C.J., Berrios, J., Colvin, J.S., Bousquet-Moore, D., et al. (2011). Synaptic dysfunction and abnormal behaviors in mice lacking major isoforms of Shank3. Hum. Mol. Genet. 20, 3093–3108.

18. Peixoto, R.T., Wang, W., Croney, D.M., Kozorovitskiy, Y., and Sabatini, B.L. (2016). Early hyperactivity and precocious maturation of corticostriatal circuits in Shank3B(−/−) mice. Nat. Neurosci. 19, 716–724.

19. Pagani, M., Bertero, A., Liska, A., Galbusera, A., Sabbioni, M., Barsotti, N., Colenbier, N., Marinazzo, D., Scattoni, M.L., Pasqualetti, M., et al. (2019). Deletion of autism risk gene Shank3 disrupts prefrontal connectivity. J. Neurosci. 39, 5299–5310.

20. Chen, Q., Deister, C.A., Gao, X., Guo, B., Lynn-Jones, T., Chen, N., Wells, M.F., Liu, R., Goard, M.J., Dimidschstein, J., et al. (2020). Dysfunction of cortical GABAergic neurons leads to sensory hyper-reactivity in a Shank3 mouse model of ASD. Nat. Neurosci. 23, 520–532.

21. Balasco, L., Pagani, M., Pangrazzi, L., Chelini, G., Ciancone Chama, A.G., Shlosman, E., Mattioni, L., Galbusera, A., Iurilli, G., Provenzano, G., et al. (2022). Abnormal whisker-dependent behaviors and altered Cortico-hippocampal connectivity in Shank3b−/− mice. Cereb. Cortex 32, 3042–3056.

22. Montagni, E., Ambrosone, M., Martello, A., Curti, L., Polverini, F., Baroncelli, L., Mannaioni, G., Pavone, F.S., Masi, A., and Allegra Mascaro, A.L. (2025). Age-dependent cortical overconnectivity in Shank3 mice is reversed by anesthesia. Transl. Psychiatry 15, 154.

23. MacDowell Camden J., Tafazoli Sina, Buschman Timothy J (2022). A Goldilocks theory of cognitive control: Balancing precision and efficiency with low-dimensional control states. Curr. Opin. Syst. Biol. 76. 10.1016/j.coisb.2017.01.006.

24. MacDowell, C.J., Libby, A., Jahn, C.I., Tafazoli, S., Ardalan, A., and Buschman, T.J. (2025). Multiplexed subspaces route neural activity across brain-wide networks. Nat. Commun. 16, 3359.

25. Zhang, Q., Yao, J., Guang, Y., Liang, S., Guan, J., Qin, H., Liao, X., Jin, W., Zhang, J., Pan, J., et al. (2017). Locomotion-related population cortical Ca2+ transients in freely behaving mice. Front. Neural Circuits 11, 24.

26. Ren, C., and Komiyama, T. (2021). Characterizing cortex-wide dynamics with wide-field calcium imaging. J. Neurosci. 41, 4160–4168.

27. Montagni, E., Resta, F., Tort-Colet, N., Scaglione, A., Mazzamuto, G., Destexhe, A., Pavone, F.S., and Allegra Mascaro, A.L. (2024). Mapping brain state-dependent sensory responses across the mouse cortex. iScience 27, 109692.

28. Benisty, H., Barson, D., Moberly, A.H., Lohani, S., Tang, L., Coifman, R.R., Crair, M.C., Mishne, G., Cardin, J.A., and Higley, M.J. (2024). Rapid fluctuations in functional connectivity of cortical networks encode spontaneous behavior. Nat. Neurosci. 27, 148–158.

29. Quarta, E., Scaglione, A., Lucchesi, J., Sacconi, L., Allegra Mascaro, A.L., and Pavone, F.S. (2022). Distributed and localized dynamics emerge in the mouse neocortex during reach-to-grasp behavior. J. Neurosci. 42, 777–788.

30. Montagni, E., Resta, F., Conti, E., Scaglione, A., Pasquini, M., Micera, S., Mascaro, A.L.A., and Pavone, F.S. (2019). Wide-field imaging of cortical neuronal activity with red-shifted functional indicators during motor task execution. J. Phys. D Appl. Phys. 52, 074001.

31. Scaglione, A., Conti, E., Allegra Mascaro, A.L., and Pavone, F.S. (2022). Tracking the effect of therapy with single-trial based classification after stroke. Front. Syst. Neurosci. 16, 840922.

32. Cecchini, G., Scaglione, A., Allegra Mascaro, A.L., Checcucci, C., Conti, E., Adam, I., Fanelli, D., Livi, R., Pavone, F.S., and Kreuz, T. (2021). Cortical propagation tracks functional recovery after stroke. PLoS Comput. Biol. 17, e1008963.

33. MacDowell, C.J., and Buschman, T.J. (2020). Low-dimensional spatiotemporal dynamics underlie cortex-wide neural activity. Curr. Biol. 30, 2665–2680.e8.

34. Mohajerani, M.H., Chan, A.W., Mohsenvand, M., LeDue, J., Liu, R., McVea, D.A., Boyd, J.D., Wang, Y.T., Reimers, M., and Murphy, T.H. (2013). Spontaneous cortical activity alternates between motifs defined by regional axonal projections. Nat. Neurosci. 16, 1426–1435.

35. Linden, N.J., Tabuena, D.R., Steinmetz, N.A., Moody, W.J., Brunton, S.L., and Brunton, B.W. (2021). Go with the FLOW: visualizing spatiotemporal dynamics in optical widefield calcium imaging. J. R. Soc. Interface 18, 20210523.

36. Nakai, N., Sato, M., Yamashita, O., Sekine, Y., Fu, X., Nakai, J., Zalesky, A., and Takumi, T. (2023). Virtual reality-based real-time imaging reveals abnormal cortical dynamics during behavioral transitions in a mouse model of autism. Cell Rep. 42, 112258.

37. MacDowell, C.J., Briones, B.A., Lenzi, M.J., Gustison, M.L., and Buschman, T.J. (2024). Differences in the expression of cortex-wide neural dynamics are related to behavioral phenotype. Curr. Biol. 34, 1333–1340.e6.

38. Michelson, N.J., Bolaños, F., Bolaños, L.A., Balbi, M., LeDue, J.M., and Murphy, T.H. (2023). Meso-py: Dual brain cortical calcium imaging in mice during head-fixed social stimulus presentation. eNeuro 10. 10.1523/ENEURO.0096-23.2023.

39. Stringer, C., Pachitariu, M., Steinmetz, N., Reddy, C.B., Carandini, M., and Harris, K.D. (2019). Spontaneous behaviors drive multidimensional, brainwide activity. Science 364, 255.

40. Hasnain, M.A., Birnbaum, J.E., Ugarte Nunez, J.L., Hartman, E.K., Chandrasekaran, C., and Economo, M.N. (2025). Separating cognitive and motor processes in the behaving mouse. Nat. Neurosci. 28, 640–653.

41. Lara, A.H., Elsayed, G.F., Zimnik, A.J., Cunningham, J.P., and Churchland, M.M. (2018). Abstract. Preprint at eLife Sciences Publications, Ltd, 10.7554/elife.31826.001 10.7554/elife.31826.001.

42. Matas, E., Maisterrena, A., Thabault, M., Balado, E., Francheteau, M., Balbous, A., Galvan, L., and Jaber, M. (2021). Major motor and gait deficits with sexual dimorphism in a Shank3 mutant mouse model. Mol. Autism 12, 2.

43. Bauer, H.F., Delling, J.P., Bockmann, J., Boeckers, T.M., and Schön, M. (2022). Development of sex- and genotype-specific behavioral phenotypes in a Shank3 mouse model for neurodevelopmental disorders. Front. Behav. Neurosci. 16, 1051175.

44. Mackevicius, E.L., Bahle, A.H., Williams, A.H., Gu, S., Denisenko, N.I., Goldman, M.S., and Fee, M.S. (2019). Abstract. Preprint at eLife Sciences Publications, Ltd, 10.7554/elife.38471.001 10.7554/elife.38471.001.

45. Vann, S.D., Aggleton, J.P., and Maguire, E.A. (2009). What does the retrosplenial cortex do? Nat. Rev. Neurosci. 10, 792–802.

46. Lyamzin, D., and Benucci, A. (2019). The mouse posterior parietal cortex: Anatomy and functions. Neurosci. Res. 140, 14–22.

47. Stacho, M., and Manahan-Vaughan, D. (2022). Mechanistic flexibility of the retrosplenial cortex enables its contribution to spatial cognition. Trends Neurosci. 45, 284–296.

48. Cui, H., and Andersen, R.A. (2007). Posterior parietal cortex encodes autonomously selected motor plans. Neuron 56, 552–559.

49. Franco, L.M., and Goard, M.J. (2021). A distributed circuit for associating environmental context with motor choice in retrosplenial cortex. Sci. Adv. 7, eabf9815.

50. Li, N., Chen, T.-W., Guo, Z.V., Gerfen, C.R., and Svoboda, K. (2015). A motor cortex circuit for motor planning and movement. Nature 519, 51–56.

51. Yamawaki, N., Radulovic, J., and Shepherd, G.M.G. (2016). A corticocortical circuit directly links retrosplenial cortex to M2 in the mouse. J. Neurosci. 36, 9365–9374.

52. Lim, D.H., Mohajerani, M.H., Ledue, J., Boyd, J., Chen, S., and Murphy, T.H. (2012). In vivo large-scale cortical mapping using channelrhodopsin-2 stimulation in transgenic mice reveals asymmetric and reciprocal relationships between cortical areas. Front. Neural Circuits 6, 11.

53. Harrison, T.C., Ayling, O.G.S., and Murphy, T.H. (2012). Distinct cortical circuit mechanisms for complex forelimb movement and motor map topography. Neuron 74, 397–409.

54. Ambrosone, M., Montagni, E., Resta, F., Ulivi, T., Curti, L., Polverini, F., Mazzamuto, G., Mannaioni, G., Masi, A., Pavone, F.S., et al. (2025). All-optical mapping reveals distributed suppression of cortical sensory responses after optogenetic silencing. Brain Stimul. 18, 1514–1522.

55. Resta, F., Montagni, E., de Vito, G., Scaglione, A., Allegra Mascaro, A.L., and Pavone, F.S. (2022). Large-scale all-optical dissection of motor cortex connectivity shows a segregated organization of mouse forelimb representations. Cell Rep. 41, 111627.

56. Conti, E., Allegra Mascaro, A.L., and Pavone, F.S. (2019). Large scale double-path illumination system with split field of view for the all-optical study of inter-and intra-hemispheric functional connectivity on mice. Methods Protoc. 2, 11.

57. Mathis, A., Mamidanna, P., Cury, K.M., Abe, T., Murthy, V.N., Mathis, M.W., and Bethge, M. (2018). DeepLabCut: markerless pose estimation of user-defined body parts with deep learning. Nat. Neurosci. 21, 1281–1289.

58. Scott, B.B., Thiberge, S.Y., Guo, C., Tervo, D.G.R., Brody, C.D., Karpova, A.Y., and Tank, D.W. (2018). Imaging cortical dynamics in GCaMP transgenic rats with a head-mounted Widefield macroscope. Neuron 100, 1045–1058.e5.

59. Nietz, A.K., Popa, L.S., Streng, M.L., Carter, R.E., Kodandaramaiah, S.B., and Ebner, T.J. (2022). Wide-field calcium imaging of neuronal network dynamics in vivo. Biology (Basel) 11, 1601.

60. Brier, L.M., Zhang, X., Bice, A.R., Gaines, S.H., Landsness, E.C., Lee, J.-M., Anastasio, M.A., and Culver, J.P. (2022). A multivariate functional connectivity approach to mapping brain networks and imputing neural activity in mice. Cereb. Cortex 32, 1593–1607.

61. Stern, M., Shea-Brown, E., and Witten, D. (2020). Inferring the spiking rate of a population of neurons from wide-field calcium imaging. bioRxiv. 10.1101/2020.02.01.930040.

62. Levine, J.H., Simonds, E.F., Bendall, S.C., Davis, K.L., Amir, E.-A.D., Tadmor, M.D., Litvin, O., Fienberg, H.G., Jager, A., Zunder, E.R., et al. (2015). Data-driven phenotypic dissection of AML reveals progenitor-like cells that correlate with prognosis. Cell 162, 184–197.

